# Predicting Chemical Shifts with Graph Neural Networks

**DOI:** 10.1101/2020.08.26.267971

**Authors:** Ziyue Yang, Maghesree Chakraborty, Andrew D White

## Abstract

Inferring molecular structure from NMR measurements requires an accurate forward model that can predict chemical shifts from 3D structure. Current forward models are limited to specific molecules like proteins and state of the art models are not differentiable. Thus they cannot be used with gradient methods like biased molecular dynamics. Here we use graph neural networks (GNNs) for NMR chemical shift prediction. Our GNN can model chemical shifts accurately and capture important phenomena like hydrogen bonding induced downfield shift between multiple proteins, secondary structure effects, and predict shifts of organic molecules. Previous empirical NMR models of protein NMR have relied on careful feature engineering with domain expertise. These GNNs are trained from data alone with no feature engineering yet are as accurate and can work on arbitrary molecular structures. The models are also efficient, able to compute one million chemical shifts in about 5 seconds. This work enables a new category of NMR models that have multiple interacting types of macromolecules.

## 1 Introduction

NMR chemical shifts of a molecule provide detailed structural information without the sample preparation requirements of X-ray crystallography^[1]^. This means that NMR can provide detail at room temperature, at reasonable concentrations, in a physiologically relevant ensemble of conformations, and even in situ ^[2,3]^. Thus there is continued interest in methods to resolve protein structure from NMR. A key step in this process is being able to predict the NMR chemical shifts from molecular structure in a *forward model*. The forward model is used for inference to compute the statistical ensemble of structure that contribute to the observed NMRs chemical shifts in experiment. In this work, we find that graph neural networks (GNNs) have good properties as a forward model and expand the types of molecular structures that can be resolved. The process of inferring the conformational ensemble with the forward model can be done via experiment directed simulation^[4,5]^, metadynamics meta-inference^[6]^, targeted metadynamics^[7,8]^, Monte Carlo/optimization^[9,10]^, biasing with restraints^[11,12]^, Bayesian ensemble refinement^[13]^, or other simulation-based inference methods^[14–16]^. A direct method like a generative model that outputs structure directly would be preferred^[17,18]^, but a forward model that can connect the chemical shift to structure would still be part of this training.

An ideal NMR chemical shift predictor should be translationally and rotationally invariant, be sensitive to both chemically bonded and non-bonded interactions, be able to handle thousands of atoms, predict shifts for multiple atom types, and be differentiable which is required for most of the inference methods mentioned above. There are two broad classes of deep learning architectures that might satisfy these requirements: 3D point cloud neural networks methods that have these equivarianaces built-in^[19,20]^, GNNs^[21–23]†^. The conceptual difference between these two approaches are that the 3D point cloud networks first build the local environment of each atom to compute atom features and then operate and pool the atom features without considering the molecular graph, whereas the graph neural networks compute atom features using the molecular graph at each layer. Here we use graph neural networks for two reasons. The first is their flexibility of how molecular graphs can be specified: with or without distances, with or without covalent bonds, and as a sparse graph. The second reason is that the goal is to use this model in molecular simulation, where the sparse molecular graph (i.e., a neighbor list) is available as input.

GNNs are now a common approach for deep learning with molecules due to their intuitive connection to molecular graphs and good performance^[24]^. Early examples of graph neural networks can be found in Sperduti and Starita^[25]^, Scarselli *et al*. ^[26]^, Gori *et al*. ^[27]^ and recent surveys can be found in Bronstein *et al*. ^[21]^, Dwivedi *et al*. ^[22]^, Wu *et al*. ^[28]^, Battaglia *et al*. ^[29]^. The unifying idea of a “graph” neural network is that its input is a graph and its output is insensitive to the order of nodes and edges (node/edge label invariant). Although not necessarily true, the outputs are typically node features, edge features, or a graph features. Battaglia *et al*. ^[29]^ went further and has proposed a unifying notation that encompasses all graph neural networks as a series of node, edge, and graph feature operations. Unlike convolutional layers in traditional deep learning^[30]^, there are still numerous competing ideas about GNNs. Wu *et al*. ^[28]^ tested about 20 GNN across seven tasks, including chemistry datasets and found no consistently best type. They did find that message-passing methods^[31]^ worked well with other deep-learning layers and building blocks.

GNNs are being widely applied in chemistry, especially in quantum machine learning^[24,31–33]^. In this work, we have chosen message passing GNNS due to their similarity to other deep learning layers^[28]^, simplicity, and good performance^[24,28]^. Our models take the molecular graph as input where the features are the atom identities and the edges are feature vectors encoding the edge type (covalent bond or nearby neighbor) and distance. The output is the predicted NMR chemical shift for C, N, or H atoms. This approach is sometimes referred to as *enn-s2s* ^[23,34]^. Our model is trained with three datasets: the RefDB dataset of cross-referenced protein structures with NMR chemical shifts^[35]^, the SHIFTX dataset^[36]^, and a database of organic molecules^[37]^.

There are numerous existing NMR chemical shift prediction models. We first review those which are for protein structure. ProShift is a dense neural network with one hidden layer that uses 350 expert chosen input features like electronegativity or dihedral angle with neighbors^[38]^. SPARTA+ uses dense neural networks with 113 expert-chosen input features^[39]^. ShiftX+ uses an ensemble approach with boosting and uses 97 expert-chosen input features^[36]^. ShiftX2 combines ShiftX+ with homology data with a database of known proteins with chemical shift. Note that ProShift, SPARTA+, ShiftX+ and ShiftX2 are not (easily) differentiable with respect to atom positions due to the use of input features and homology data. They are also restricted to proteins due to the use of protein-specific features that are not defined for general molecules. CamShift uses a polynomial expansion of the pair-wise distances between an atom and its neighbors to approximate the NMR chemical shift^[40]^ and thus is differentiable. This has made it a popular choice^[41–43]^ and it is implemented in the PLUMED plugin^[44]^. However, CamShift does not treat side-chains and is insensitive to effects like hydrogen bonding. Of these select methods discussed, ShifX2 is typically viewed as most accurate and CamShift as the most useful for use in inferring protein structure in a molecular simulation. Our goal is to combine the high-accuracy approach of methods like ShiftX2 with the differentiable nature of CamShift. Furthermore, our approach does not require hand-engineered features and instead uses only the elements of the atoms and distances as input. This enables it to be used on both ligands and proteins.

Outside of protein structure, NMR prediction is a classic machine learning problem in chemistry. Paruzzo *et al*. ^[45]^ developed a Gaussian process regression framework for prediction of NMR chemical shifts for solids. They used smooth overlap of atomic positions (SOAP) kernel to represent the molecular structural environment. Rupp *et al*. ^[46]^ used kernel learning methods to predict chemical shfits from a small molecule training set with DFT shifts. Jonas and Kuhn^[47]^ used graph convolution neural network to predict ^1^H and ^13^C chemical shifts along with the uncertainties. Kang *et al*. ^[48]^ did similar work, again with a GNN and message passing. This is probably the most similar to our message passing GNN, but they considered small molecules and not 3D structure. Examples of others work using message passing GNNs in chemistry include Raza *et al*. ^[49]^ who predicted partial charges of metal organic frameworks, the original message passing paper byGilmer *et al*. ^[31]^ which predicted energies of molecules, and St. John *et al*. ^[50]^ who predicted bond disassociation energies. There are also first-principles methods for computing NMR chemical shifts, however we do not compare with these since their computational speed and accuracy are not comparable with empirical methods^[51–53]^.

## 2 Data Preparation

Our model was trained with three datasets. The first is a paired dataset of 2,405 proteins with both X-ray resolved crystal structures and measured NMR chemical shifts created by Zhang *et al*. ^[35]^. This was segmented into a fragment dataset of 131,015 256 atom fragments with approximately 1.25 million NMR chemical shifts. To prepare the fragments, each residue in each protein was converted into a fragment. All atoms in prior and subsequent residues were included along with residues which had an atom spatially close to the center residue, but their labels (chemical shifts) were not included. Residue *i* is close to residue *j* if an atom from residue i is one of the 16 closest non-bonded atoms of an atom in residue *j* (i.e., they share a neighbor). We did not use distance cutoffs because neighbor lists are used in subsequent stages and if an atom is not on the neighbor list, it need not be included in the fragment. Additional preprocessing was omitting fragments with missing residues, fixing missing atoms, removing solvent/heteroatoms, ensuring the NMR chemical shifts sequenced aligned with the X-ray structures, and matching chains. This was done with PDBFixer, a part of the OpenMM framework^[54]^. About 5% of residues were excluded due to these constraints and 0.93% were excluded because the resulting fragments could not fit into the 256 atom fragment. Some X-ray resolved crystal structures have multiple possible structures. We randomly sampled 3 of these (with replacement) so that some fragments may be duplicated. The number of fragments including these possible duplicates is 393,045. This dataset will be called RefDB dataset.

The second dataset was prepared identically and contains 197 in the training and 62 proteins in test. It is the SHIFTX dataset and contains 21,878 fragments for training^[36]^. This dataset is higher-quality (see training curves results) due to careful processing by Han *et al*. ^[36]^ and does not have multiple possible structures. The SHIFTX test dataset of 62 proteins (7494 fragments) was used for calculation of all test data and was not included in training. These PDB IDs were also removed from the RefDB dataset so that they did not inadvertently enter training. These protein datasets contain C, N and H chemical shifts.

The third dataset was 369 “metabolites” (biologically relevant organic molecules) from the human metabolome 4.0 database^[37]^. These were converted into 3D conformers with RDKit using the method of Riniker and Landrum^[55]^. Here, each molecule is a fragment and no segmenting of molecules was done. This is referred to as the metabolome dataset.

Each molecular fragment is 256 atoms represented as integers indicating element and each atom has up to 16 edges that connect it to both spatial and covalent neighbors. The edges contain two numbers: an encoding of the type of edge (covalent or spatial) and the distance. These two items encode the molecular graph. An example of a fragment from RefDB dataset is shown in Figure 1. This approach of using covalent bonds and spatial neighbors is somewhat analogous to attention, which is an open area of research in GNNs because its effect is not always positive^[56]^.

**Figure 1:**
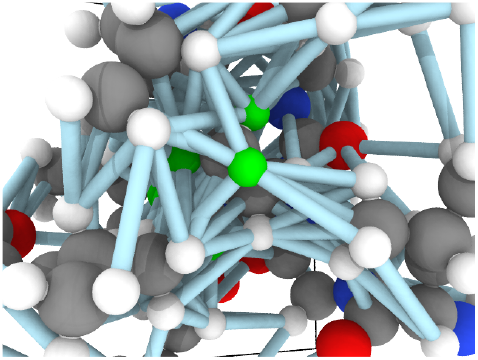
An example graph used as input to the GNN. The atoms in greens will have their chemical shifts predicted and are connected to neighboring atoms by edges, which includes both bonded and non-bonded edges. The edges are encoded as feature vectors which contains both an embedding representing the type of edge (e.g., covalent) and distance.

## 3 Model

Our GNN consists of 3 parts: (i) a dense network *ℱ* (**B**) = **N** whose input is the rank 3 (omitting batch rank) edge tensor **B** and output is the rank 3 neighbor tensor **N**; (ii) a message passing neural network *𝒢* (**X**^**0**^, **N**) whose input is the rank 2 tensor **X**^**0**^ and **N**. Its output is a rank 2 tensor **X**^*k*^; (iii) a dense network ℋ (*X*^*k*^) whose output is the chemical shifts. The architecture is shown in Figure 2. Hyperparameters were optimized on a 20/80 validation/train split of the ShiftX training dataset. The hyperparamers were layer number (1-6 explored), **X**/**N** feature dimensions (16-256, 1-32 respectively), L2 regularization^[30]^, dropout^[57]^, residue^[58]^, and the use of Schütt *et al*. ^[23]^ continuous radial basis convolutions on distance (or distance binning), choice of loss, and the use of non-linear activation in final layers. L2 regularization and dropout were found to be comparable to early-stopping on validation, so early-stop was used instead. Model training was found to diverge without residue connections, which others have seen^[59]^. Final layer numbers are *K* = 4, *L* = 3, *J* = 3. The neighbor (ℱ(**B**)) feature dimension is 4 and atom feature dimension is 256. Embedding are used for inputs. Edges use a 1D embedding for type and distance was tiled 31 times to make a 32 size input. Binning these distances seemed to have negligible affect on performance. The atom element identities were converted to a tensor with 256D embedding look-up.

**Figure 2:**
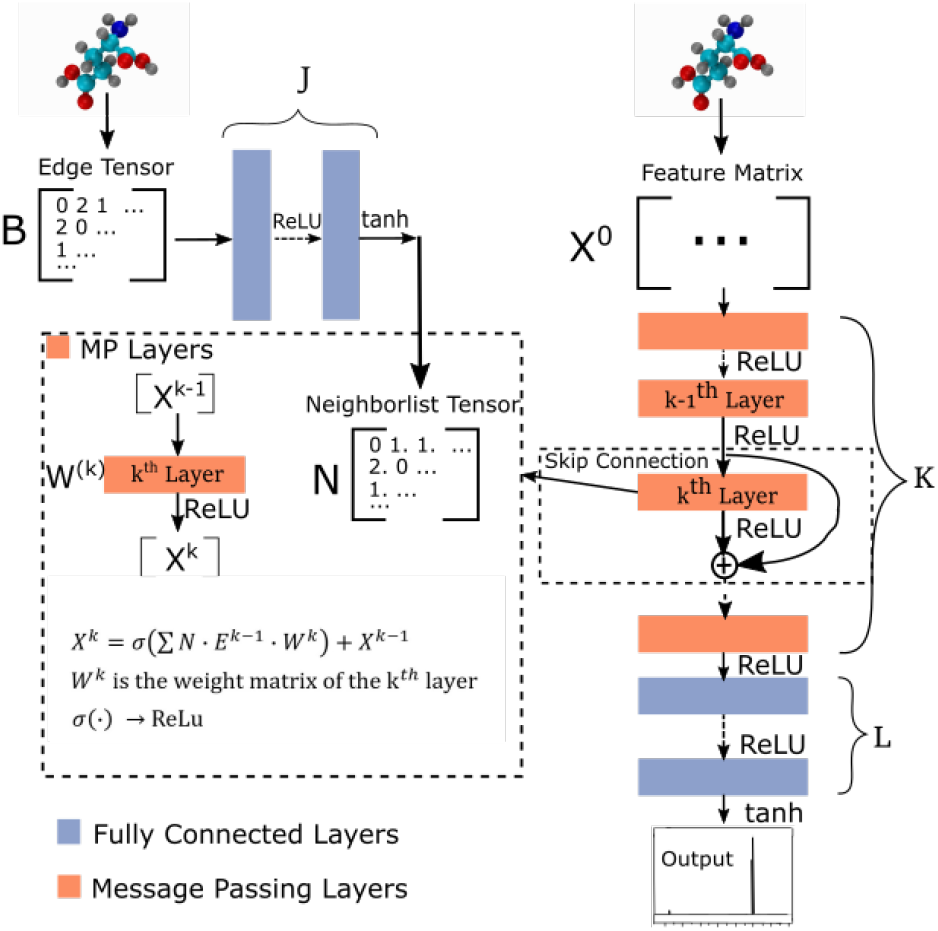
Graph Neural network architecture. **B** is input molecular graph edge features which is distance and chemical bond type (covalent or non-bonded). *N* is the output neighbor features tensor used for MP layers. *X*^0^ is input feature matrix, consisting only of element types. MP layers have residue connections are defined in Equation 1. There are K MP layers and L output FC layers. Output is passed through Equation 2 to account for element NMR differences.

*ℱ* uses ReLU activation^[60]^ except in the last layer, where tanh is used. In **G**, our message passing update function is matrix multiplication with activation applied after sum pooling across neighbors:

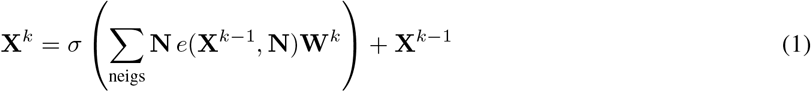

where *e*(**X**^*k*−1^, **N**) gives a rank 3 tensor **E**^*k*−2^that is the features of neighbors of each atom. *σ* is the ReLU activation function. **W**^*k*^ is rank 3 because each edge feature affects the input/output atom features separately. Instead of including an atom in its own neighbors, we use a residue connection. Our choice of message passing and lack of node update function (e.g,. GRUs in Gilmer *et al*. ^[31]^) makes it one of the simplest message passing variants. *ℋ* uses a tanh in the second to last layer and the last layer used linear activation and output dimension *Z. Z* is the number of unique elements in the dataset. Both *ℱ* and *ℋ* had bias.

Output chemical shifts 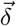 are computed as

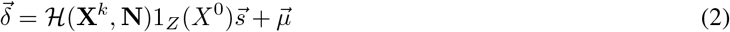

where 1_*Z*_(*X*^0^) is a one-hot indicator for atom element with *Z* columns, 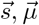, are *Z* pre-computed standard deviation and means of the refDB chemical shifts for each element. This chosen done to make labels be approximately from − 1 to 1 for training. This also has the effect of making any chemical shift for a non-trained element (e.g., N) be 0. The loss function is combined correlation and root mean squared deviation (RMSD):

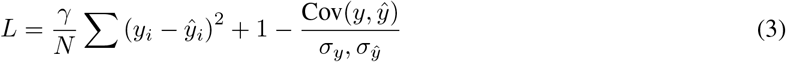

where *γ* = 0.001 for models trained on H only and 0.01 for models trained on all data. Training on correlation in addition to RMSD was found to improve model correlation. The +1 is to prevent loss from being negative and has no effect on gradients.

## 4 Training

Training was done in the TensorFlow framework^[61]^. Variables were initialized with the Glorot initializer^[62]^ and optimized with Adam optimizer^[63]^ with a learning rate schedule of [10^−3^, 10^−3^, 10^−4^, 10^−5^| 10^−4^, 10^−5^, 10^−5^| 10^−5^] where | indicates a switch to a new dataset, except the last which was joint training (see below). Early stopping with patience 5 was done for training. The first dataset was trained with 5 epochs, the second with 50, and the third was combined with the second for final training again with 50 epochs. The second and third dataset when combined have large class imbalance so rejection sampling was used at the residue level where metabolites were counted as a residue. Therefore, each amino acid and metabolites were seen with equal probability. Each epoch was one complete iteration through the dataset. Batch size was 16 fragments (16 × 256 atoms). Training and inference were found to take about 0.0015 seconds per fragment (5.7 *µ*s per shift) with the full model on a single Tesla V100 GPU. Timing was averaged on the SHIFTX dataset (21,878 fragments) with loading times excluded.

## 5 GNN Results

Unless indicated, models were trained only on H chemical shifts for assessing features and training curves. Training on all types requires the metabolome dataset and more complex joint training with rejection sampling. A log-log training curve is shown in Figure 3 which shows H^*α*^ accuracy on the SHIFTX test dataset as a function of amount of training data. 100% here means all trainnig data exlcuding validation. The SHIFTX dataset is about one tenth the size of RefDB dataset but can provide nearly the same accuracy as shown (0.29 vs 0.26 RMSD). The RefDB dataset and SHIFTX dataset contain the same proteins, but the SHIFTX dataset are more carefully processed. This shows more careful processing of data is more important than number of structures.

**Figure 3:**
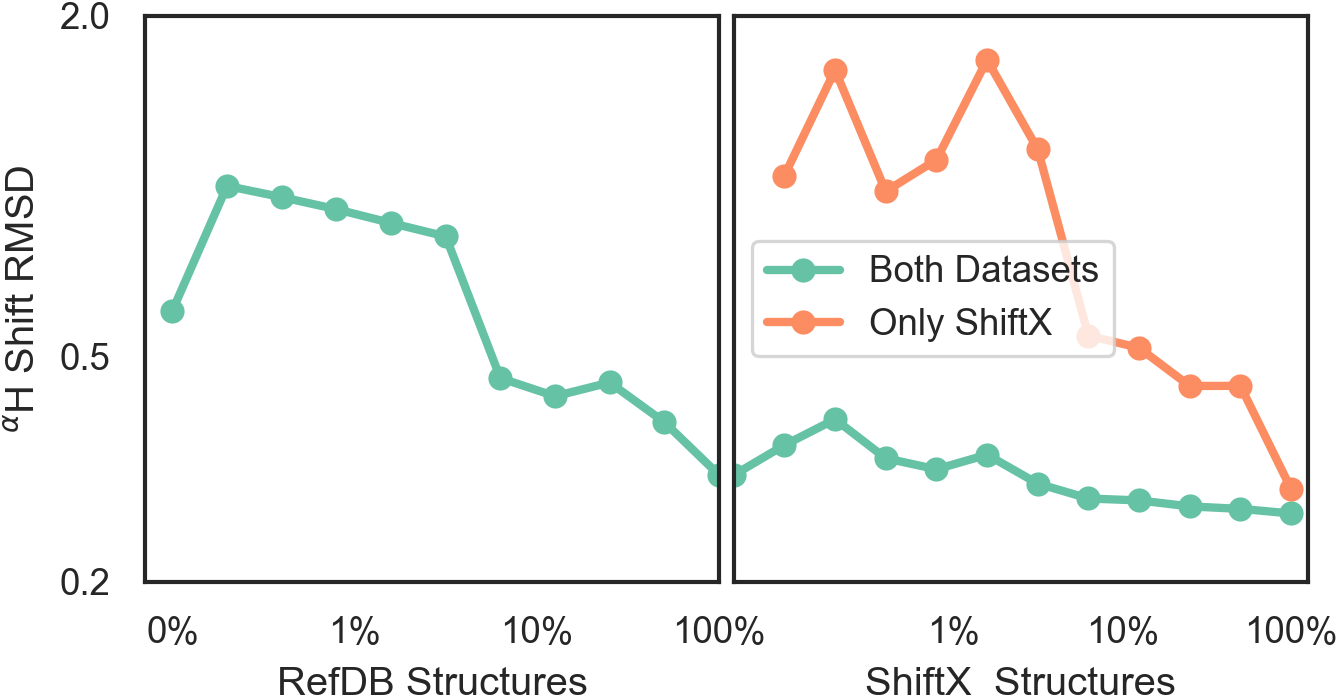
A log-log plot of training root mean squared deviation of labels with model predicted chemical shift of H^*α*^ as a function of elements in dataset. 100% means all data excluding validation and test data is provided. The number of RefDB dataset examples is 131,015 (716,164 shifts) and SHIFTX dataset is 21,878 examples (88,392 shifts).

The final model performance with all training data is shown in Table 1. A complete breakdown per amino acid and atom name for all models is given in Supporting Information. Comparisons were done using the SHIFTX+ webserver^†^ and the latest implementation of CS2Backbone in Plumed^[44]^. We also include the reported performance of SHIFTX+ on their website that had better performance, which could be because in our training and comparisons we did not set pH and temperatures and instead used pH = 5, temperature = 298K. Our rationale for this decision was that we wanted a model whose input is *only* molecular structure, and not experimental details such as buffer, pH, temperature etc. Thus we compared to other models with the same restriction. Overall both the model with H shift only and all elements perform comparably with existing methods but the GNN is not state of the art. The advantage of our model is the efficiency and ability to input any molecule. Table 1 also shows the effect of changing parameter number. There seems to be a sharp transition at the million parameters, meaning models that are much smaller can be used for intermediate accuracy. Some of the major choices of architecture design are also shown: including using dropout (in *ℱ G ℋ*), example weighting by class (amino acid), and without non-linear activation. The label variance is computed by comparing repeat measurements of the same protein structure in the RefDB dataset and should be taken as the upper-limit beyond which experimental error is more important. This non-linear scaling of accuracy with parameter number has been previously observed in GNNs^[64]^.

**Table 1:**
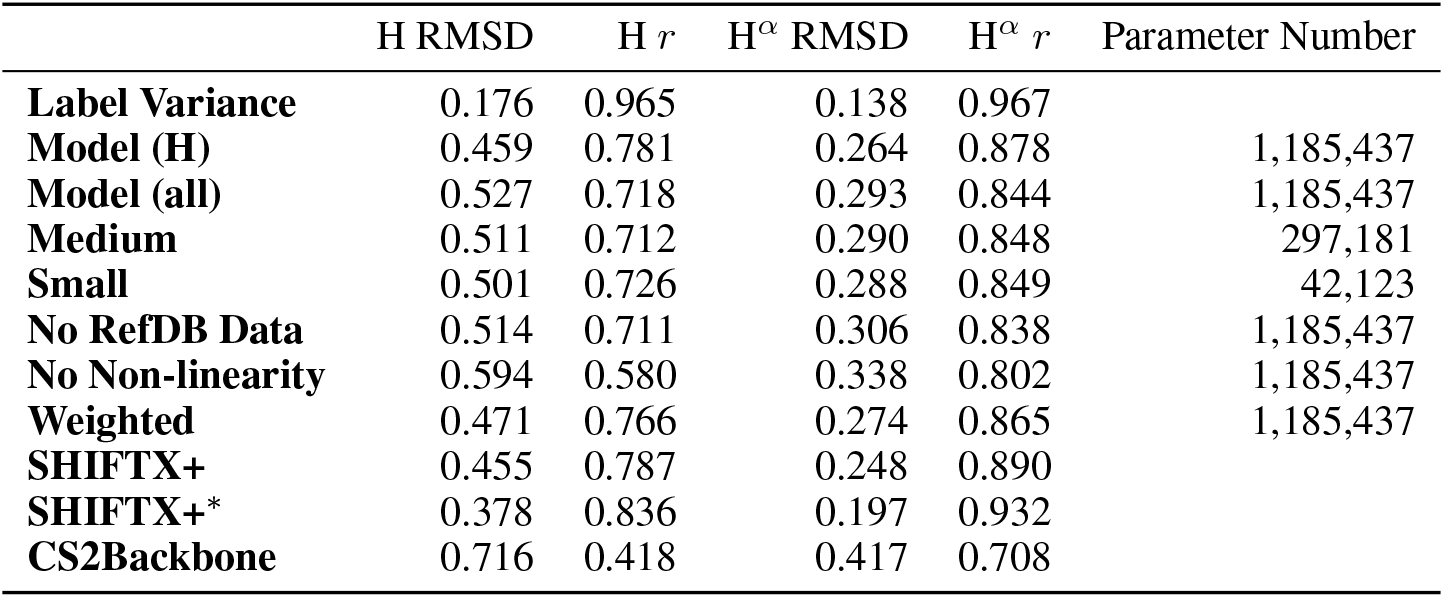
A comparison of the GNN presented here, other similar NMR models, and how model size affects performance. ^*^Reported by SHIFTX+ developers, which includes temperature and pH effects. All others were computed independently in this work.

| | H RMSD | H $r$ | H$^{\alpha}$ RMSD | H$^{\alpha}$ $r$ | Parameter Number |
| --- | --- | --- | --- | --- | --- |
| **Label Variance** | 0.176 | 0.965 | 0.138 | 0.967 |  |
| **Model (H)** | 0.459 | 0.781 | 0.264 | 0.878 | 1,185,437 |
| **Model (all)** | 0.527 | 0.718 | 0.293 | 0.844 | 1,185,437 |
| **Medium** | 0.511 | 0.712 | 0.290 | 0.848 | 297,181 |
| **Small** | 0.501 | 0.726 | 0.288 | 0.849 | 42,123 |
| **No RefDB Data** | 0.514 | 0.711 | 0.306 | 0.838 | 1,185,437 |
| **No Non-linearity** | 0.594 | 0.580 | 0.338 | 0.802 | 1,185,437 |
| **Weighted** | 0.471 | 0.766 | 0.274 | 0.865 | 1,185,437 |
| **SHIFTX+** | 0.455 | 0.787 | 0.248 | 0.890 |  |
| **SHIFTX+\*** | 0.378 | 0.836 | 0.197 | 0.932 |  |
| **CS2Backbone** | 0.716 | 0.418 | 0.417 | 0.708 |  |

Figure 4 shows the effect of input features on the model. Model performance is good with a normal molecular graph GNN (*No Distances*) where the input is only which atoms are near and which are covalently bonded. Knowing the distance provides a small improvement in accuracy. Knowing which atoms are spatially near provides a larger improvement, as shown in the *Only Chemical Bonded* model. None of the models are close to the *Label Variance*, which is the upper-bound of what is possible.

**Figure 4:**
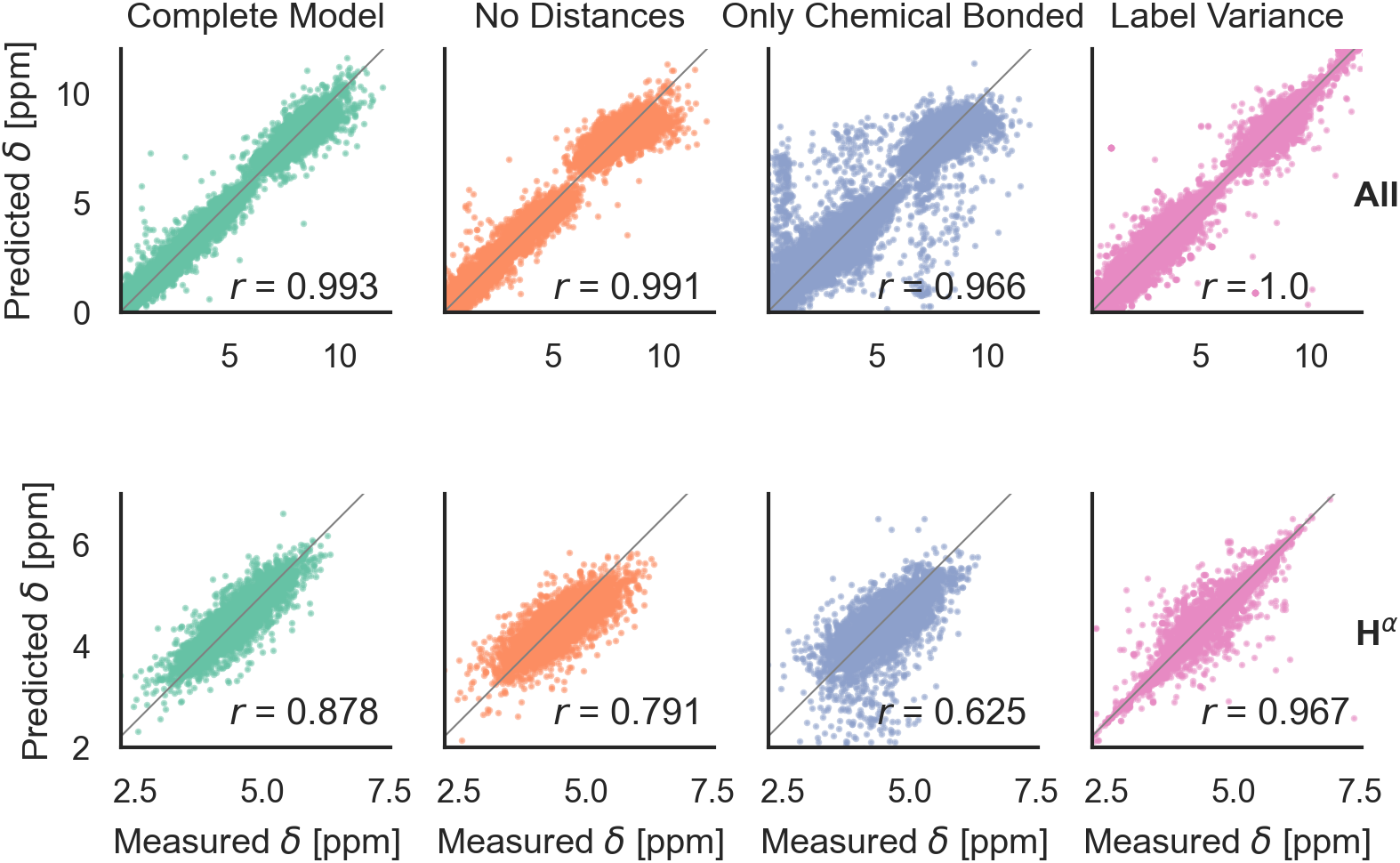
Parity plots comparing edge features in the GNN. *No Distances* means that non-bonded neighbors are included, but with no distances. *Only Chemical Bonded* means distance is included but only neighbors directly covalently bonded with an atom are included. *Label Variance* is the variation between repeat measured NMR chemical shifts in the RefDB dataset^[35]^ and should be taken as the upper-limit beyond which experimental errors are more significant than model fit.

## 6 Multitype Model

After training on all element types and with metabolome dataset, model accuracy decreased slightly (Table 1). However, the model has the desired features as shown in Figure 5. It is able to model C, N, and H chemical shifts with good correlation and good RMSDs (N: 2.982, C: 1.652, 0.368). The correlation on the important H^*α*^ is 0.844 vs 0.878 in the H model. Including metabolome dataset into training gives a 0.872 correlation on the withheld 20% test (74 molecules). No validation was used for this data because hyperparameters were not tuned. Training on *only* metabolome dataset gives 0.92 correlation on withheld data and could be taken as an approximate upper-bound because the ratio of trainable parameters (1 million) to data (369) is extreme.

**Figure 5:**
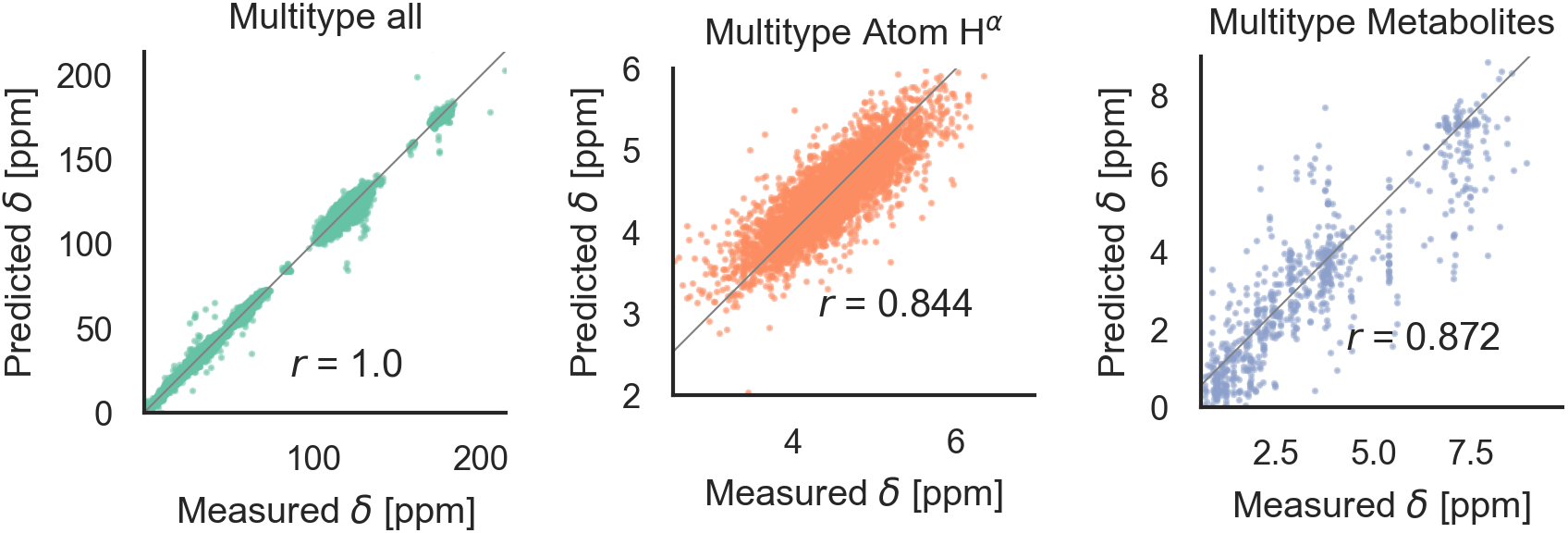
Parity plots for the multitype model, which can treat C, N, H atoms and organic molecules. *Multitype all* is the combined plot for C, N, and H in test proteins. *Multitype Atom H*^*α*^ shows the performance of this model on the important H^*α*^ atom type. *Metabolites* is the model performance on metabolites^[37]^.

Figure 6 shows phenomenological validation of the GNN model on two untrained properties: sensitivity to secondary structure and chemical shift in hydrogen bonding. The left panel shows the average predicted chemical shifts of each amino acid and secondary structure combination. As expected based on model performance, it does well at predicting the effect of secondary structure on chemical shift. Disagreement is seen on less frequently observed combinations like cystein *β*-sheets and Tryptophan. Most comparable models like ProShift or ShiftX^[36,38,39]^ have secondary structure (or dihedral angles) as inputs for computing chemical shifts. The end-to-end training of the GNN captures this effect. The results are consistent with previous studies^[65–67]^ which showed downfield shift of H^*α*^ *δ* for *β*-sheet and upfield shift for *α*-helix. The right panel shows the effect of breaking a salt bridge (ionic hydrogen bond) between an arginine and glutamic acid on the H^*ϵ*^ chemical shift. This atom was chosen because it is observable in solution NMR. White *et al*. ^[68]^ computed the chemical shift change to be 0.26 Δ*δ* ppm for breaking this hydrogen bond based on single-amino acid mixture NMR. The molecular graph was fixed here to avoid effects of neighbor lists changing. The model gets a similar upfield shift and thus shows it could be used to model protein-protein interfaces where side-chain – side-chain interactions are critical. It is also consistent with previous reports^[69,70]^ where an increasing strength of hydrogen bond was associated with greater deshielding and subsequent downfield shift of H^*α*^ *δ*.

**Figure 6:**
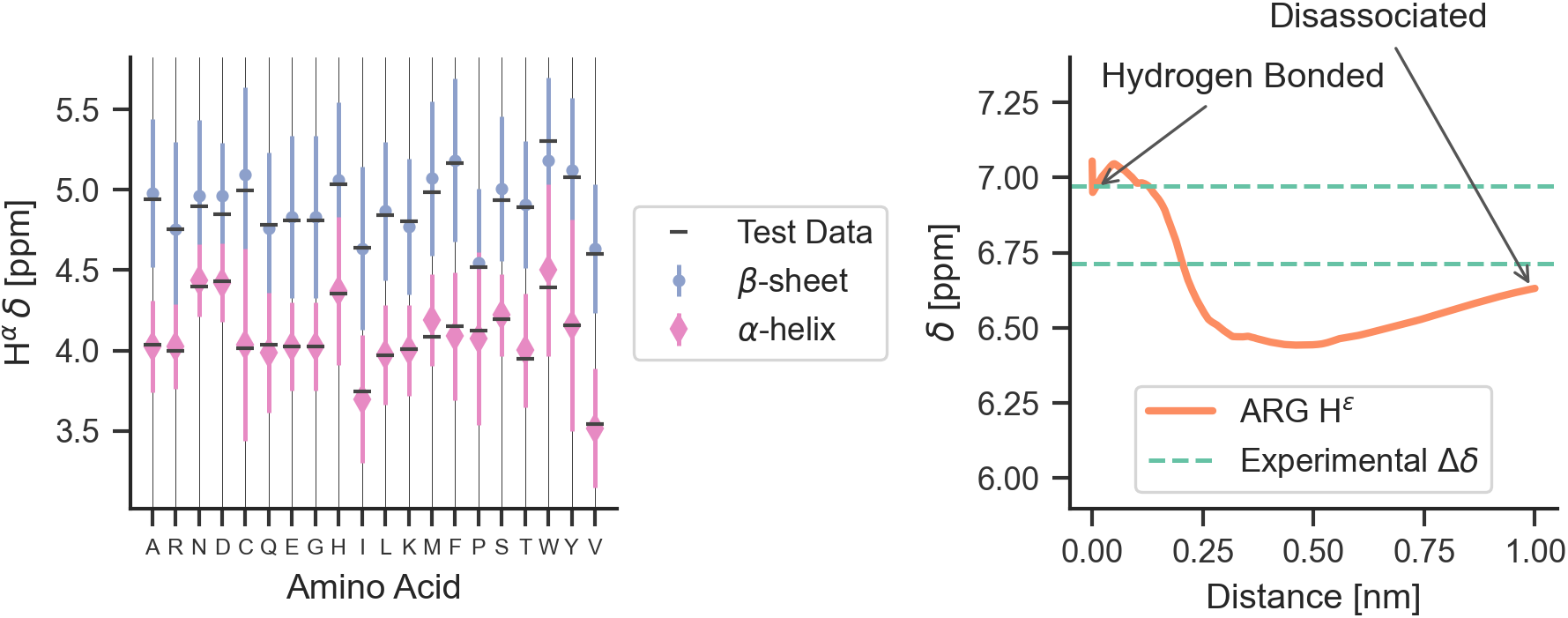
The model performance on secondary structure and inter-molecular interactions. Left panel shows the effects of secondary structure on *α* H *δ*. Each colored point is the average predicted across test data for amino acid/secondary structure combination. Vertical lines indicate uncertainty. Horizontal line indicates true average from data. Right panel shows the downfield shift of protons participating in a salt bridge (ionic hydrogen bond) between an arginine and glutamic amino acid on separate chains. Experimental data is from White *et al*. ^[68]^ indicates relative difference in chemical shift of the NH^*ϵ*^ proton between an amidated/acetylated ARG – GLU mixed solution vs amidated/acetylated ARG alone.

## 7 Discussion

The GNN is able to compute chemical shifts for arbitrary molecules, be sensitive to both covalent and non-bonded interactions, parse a million chemical shifts in 5 seconds, and is differentiable with respect to pairwise distances. Model accuracy is near state of the art performance. There is a trade-off between the chemical elements to train for and model accuracy, leaving an unanswered question of if more trainable parameters are required. Training is complex, because there are three datasets and they are of varying quality and sizes. Effort should be invested in better quality protein structure data. Finally, there is a large number of message passing choices and more exploration could be done.

## 8 Conclusion

This work presents a new class of chemical shift predictors that requires no a priori knowledge about what features affect chemical shift. The GNN input is only the underlying molecular graph and elements and requires no details about amino acids, protein secondary structure or other features. The GNN is close to state of the art in performance and able to take arbitrary input molecules, including organic molecules. The model is highly-efficient and differentiable, making it possible to use in molecular simulation. Important physical properties also arise purely from training: *β*-sheets formation causes downfield shifts and breaking salt bridges causes upfield shifts. This work opens a new direction for connecting NMR experiments to molecular structure via deep learning.

## 9 Acknowledgements

This material is based upon work supported by the National Science Foundation under Grant No. (1764415 and 1751471). We thank the Center for Integrated Research Computing(CIRC) at the University of Rochester for providing computational resources and technical support.

## Footnotes

† We do not consider featurization like computing dihedral angles or electronegativity of atoms because they cannot generalize to arbitrary structures and derivatives do not always exist.

† http://shiftx2.ca/

## Notes

### Competing Interest Statement

The authors have declared no competing interest.

