## Supporting Information for "Predicting Chemical Shifts with Graph Neural Networks"

August 26, 2020

Table S1: Complete model stats on test data across classes and atom types for Model (H).

| title | corr-coeff | R^2 | MAE | RMSD | N |
|---|---|---|---|---|---|
| overall | 0.9935 | 0.9870 | 0.2218 | 0.3235 | 32520 |
| overall-H | 0.9935 | 0.9870 | 0.2218 | 0.3235 | 32520 |
| class/ALA | 0.9867 | 0.9736 | 0.2414 | 0.3311 | 1046 |
| class/ARG | 0.9925 | 0.9850 | 0.2149 | 0.3210 | 1747 |
| class/ASN | 0.9894 | 0.9789 | 0.2575 | 0.3556 | 946 |
| class/ASP | 0.9891 | 0.9783 | 0.2489 | 0.3615 | 1430 |
| class/CYS | 0.9812 | 0.9627 | 0.3290 | 0.4789 | 291 |
| class/GLU | 0.9943 | 0.9887 | 0.1926 | 0.2793 | 2144 |
| class/GLN | 0.9925 | 0.9850 | 0.2178 | 0.3159 | 1108 |
| class/GLY | 0.9812 | 0.9627 | 0.3141 | 0.4281 | 1391 |
| class/HIS | 0.9859 | 0.9720 | 0.2923 | 0.3871 | 521 |
| class/ILE | 0.9952 | 0.9905 | 0.1974 | 0.2768 | 2917 |
| class/LEU | 0.9945 | 0.9890 | 0.2055 | 0.2803 | 4557 |
| class/LYS | 0.9938 | 0.9877 | 0.1721 | 0.2653 | 3241 |
| class/MET | 0.9905 | 0.9811 | 0.2489 | 0.3616 | 524 |
| class/PHE | 0.9846 | 0.9694 | 0.2612 | 0.3600 | 1812 |
| class/PRO | 0.9491 | 0.9009 | 0.2328 | 0.3404 | 1273 |
| class/SER | 0.9822 | 0.9647 | 0.2513 | 0.3802 | 1193 |
| class/THR | 0.9908 | 0.9817 | 0.2198 | 0.3895 | 1683 |
| class/TRP | 0.9831 | 0.9664 | 0.2897 | 0.4129 | 485 |
| class/TYR | 0.9875 | 0.9752 | 0.2448 | 0.3327 | 1047 |
| class/VAL | 0.9954 | 0.9907 | 0.1928 | 0.2744 | 3164 |
| names/H | 0.7803 | 0.6089 | 0.3389 | 0.4596 | 6859 |
| names/HA | 0.8790 | 0.7727 | 0.1935 | 0.2628 | 5031 |

Continued on next page

*

Table S1: Complete model stats on test data across classes and atom types for Model (H).

| title | corr-coeff | R^2 | MAE | RMSD | N |
|---|---|---|---|---|---|
| names/HB2 | 0.9219 | 0.8499 | 0.2120 | 0.2917 | 3105 |
| names/HB3 | 0.9207 | 0.8477 | 0.2196 | 0.3051 | 2950 |
| names/HG2 | 0.8728 | 0.7618 | 0.1580 | 0.2468 | 1202 |
| names/HG3 | 0.8902 | 0.7925 | 0.1676 | 0.2395 | 1094 |
| names/HD2 | 0.9913 | 0.9826 | 0.1912 | 0.2911 | 957 |
| names/HD3 | 0.9651 | 0.9315 | 0.1680 | 0.2476 | 578 |
| names/HB | 0.9609 | 0.9233 | 0.1993 | 0.3088 | 881 |
| names/HG12 | 0.6101 | 0.3723 | 0.1459 | 0.1954 | 323 |
| names/HG13 | 0.6807 | 0.4634 | 0.1348 | 0.1798 | 323 |
| names/HG21 | 0.7792 | 0.6072 | 0.1555 | 0.2090 | 870 |
| names/HG22 | 0.7626 | 0.5815 | 0.1593 | 0.2145 | 870 |
| names/HG23 | 0.7582 | 0.5748 | 0.1573 | 0.2162 | 870 |
| names/HD11 | 0.7086 | 0.5021 | 0.1576 | 0.2054 | 695 |
| names/HD12 | 0.6769 | 0.4582 | 0.1633 | 0.2139 | 695 |
| names/HD13 | 0.6985 | 0.4880 | 0.1569 | 0.2075 | 695 |
| names/HG11 | 0.6818 | 0.4649 | 0.1353 | 0.1799 | 323 |
| names/HE | 0.5674 | 0.3219 | 0.3463 | 0.6170 | 85 |
| names/HE2 | 0.9939 | 0.9879 | 0.1541 | 0.2190 | 519 |
| names/HE3 | 0.9904 | 0.9809 | 0.1409 | 0.2225 | 265 |
| names/HZ2 | 0.7137 | 0.5093 | 0.2223 | 0.2962 | 43 |
| names/HZ3 | 0.3677 | 0.1352 | 0.2335 | 0.3057 | 34 |
| names/HG | 0.8169 | 0.6673 | 0.2256 | 0.3578 | 348 |
| names/HD21 | 0.7159 | 0.5126 | 0.1503 | 0.1992 | 387 |
| names/HD22 | 0.7206 | 0.5193 | 0.1488 | 0.1962 | 387 |
| names/HD23 | 0.6993 | 0.4891 | 0.1566 | 0.2038 | 387 |
| names/HD1 | 0.6081 | 0.3698 | 0.2196 | 0.2995 | 351 |
| names/HE1 | 0.9349 | 0.8740 | 0.2689 | 0.4067 | 375 |
| names/HA2 | 0.5470 | 0.2992 | 0.2699 | 0.3631 | 428 |
| names/HA3 | 0.4195 | 0.1760 | 0.3237 | 0.4448 | 417 |
| names/HZ | 0.6896 | 0.4755 | 0.2259 | 0.3129 | 123 |
| names/HH2 | 0.4505 | 0.2029 | 0.1938 | 0.2504 | 40 |

Table S2: Complete model stats on test data across classes and atom types for Model (all).

| title | corr-coeff | R^2 | MAE | RMSD | N |
|---|---|---|---|---|---|
| overall | 0.9997 | 0.9994 | 0.7988 | 1.4570 | 65163 |
| overall-N | 0.9183 | 0.8432 | 2.1260 | 2.9820 | 7265 |

Continued on next page

Table S2: Complete model stats on test data across classes and atom types for Model (all).

| title | corr-coeff | R^2 | MAE | RMSD | N |
|---|---|---|---|---|---|
| overall-C | 0.9997 | 0.9993 | 1.1190 | 1.6520 | 25378 |
| overall-H | 0.9916 | 0.9832 | 0.2522 | 0.3676 | 32520 |
| class/ALA | 0.9998 | 0.9996 | 0.8593 | 1.3090 | 3307 |
| class/ARG | 0.9995 | 0.9991 | 0.7814 | 1.6900 | 3454 |
| class/ASN | 0.9997 | 0.9993 | 0.9936 | 1.5960 | 2319 |
| class/ASP | 0.9997 | 0.9994 | 0.8585 | 1.4270 | 3158 |
| class/CYS | 0.9996 | 0.9992 | 1.1620 | 1.7840 | 680 |
| class/GLU | 0.9997 | 0.9995 | 0.7212 | 1.3250 | 4368 |
| class/GLN | 0.9997 | 0.9994 | 0.8434 | 1.4370 | 2458 |
| class/GLY | 0.9997 | 0.9994 | 0.8926 | 1.4930 | 2956 |
| class/HIS | 0.9987 | 0.9974 | 1.2830 | 3.0880 | 1112 |
| class/ILE | 0.9997 | 0.9994 | 0.6935 | 1.2470 | 5415 |
| class/LEU | 0.9998 | 0.9995 | 0.6386 | 1.1040 | 7948 |
| class/LYS | 0.9998 | 0.9995 | 0.6461 | 1.1720 | 6018 |
| class/MET | 0.9996 | 0.9993 | 0.9166 | 1.5220 | 1124 |
| class/PHE | 0.9994 | 0.9989 | 1.1840 | 2.2350 | 3546 |
| class/PRO | 0.9999 | 0.9997 | 0.5865 | 0.9146 | 2463 |
| class/SER | 0.9997 | 0.9993 | 0.8792 | 1.5200 | 2683 |
| class/THR | 0.9997 | 0.9994 | 0.8144 | 1.3980 | 3412 |
| class/TRP | 0.9995 | 0.9990 | 1.1910 | 1.9550 | 982 |
| class/TYR | 0.9997 | 0.9994 | 0.9679 | 1.5110 | 2068 |
| class/VAL | 0.9998 | 0.9995 | 0.6532 | 1.1990 | 5692 |
| names/N | 0.8795 | 0.7735 | 2.1070 | 2.7540 | 6822 |
| names/C | 0.8268 | 0.6836 | 0.9708 | 1.2880 | 6012 |
| names/H | 0.7180 | 0.5156 | 0.3914 | 0.5274 | 6859 |
| names/CA | 0.9704 | 0.9418 | 0.9562 | 1.2600 | 7385 |
| names/CB | 0.9916 | 0.9832 | 1.1640 | 1.7620 | 6069 |
| names/CG | 0.9990 | 0.9980 | 0.9336 | 1.3190 | 1513 |
| names/CD | 0.9997 | 0.9993 | 0.8187 | 1.1310 | 666 |
| names/HA | 0.8442 | 0.7126 | 0.2185 | 0.2931 | 5031 |
| names/HB2 | 0.9049 | 0.8189 | 0.2320 | 0.3212 | 3105 |
| names/HB3 | 0.9012 | 0.8122 | 0.2405 | 0.3372 | 2950 |
| names/HG2 | 0.8538 | 0.7289 | 0.1823 | 0.2648 | 1202 |
| names/HG3 | 0.8619 | 0.7429 | 0.1934 | 0.2683 | 1094 |
| names/HD2 | 0.9887 | 0.9776 | 0.2185 | 0.3308 | 957 |
| names/HD3 | 0.9491 | 0.9008 | 0.2024 | 0.3000 | 578 |
| names/CG1 | 0.9065 | 0.8218 | 0.9525 | 1.4850 | 548 |
| names/CG2 | 0.8283 | 0.6861 | 1.1040 | 1.4470 | 783 |
| names/CD1 | 0.9996 | 0.9991 | 1.4960 | 1.9270 | 886 |

Continued on next page

Table S2: Complete model stats on test data across classes and atom types for Model (all).

| title | corr-coeff | R^2 | MAE | RMSD | N |
|---|---|---|---|---|---|
| names/HB | 0.9521 | 0.9066 | 0.2180 | 0.3381 | 881 |
| names/HG12 | 0.5911 | 0.3494 | 0.1584 | 0.2102 | 323 |
| names/HG13 | 0.6414 | 0.4114 | 0.1496 | 0.1990 | 323 |
| names/HG21 | 0.6602 | 0.4358 | 0.1821 | 0.2561 | 870 |
| names/HG22 | 0.6827 | 0.4661 | 0.1834 | 0.2457 | 870 |
| names/HG23 | 0.6720 | 0.4516 | 0.1804 | 0.2471 | 870 |
| names/HD11 | 0.6201 | 0.3846 | 0.1715 | 0.2323 | 695 |
| names/HD12 | 0.6111 | 0.3735 | 0.1754 | 0.2340 | 695 |
| names/HD13 | 0.6438 | 0.4145 | 0.1712 | 0.2283 | 695 |
| names/HG11 | 0.6327 | 0.4003 | 0.1536 | 0.2012 | 323 |
| names/NE | 0.3560 | 0.1267 | 2.2390 | 6.7040 | 80 |
| names/CZ | 0.9592 | 0.9200 | 3.6790 | 5.2430 | 117 |
| names/HE | 0.4747 | 0.2254 | 0.3819 | 0.6412 | 85 |
| names/CE | 0.9835 | 0.9673 | 1.1730 | 1.8420 | 309 |
| names/HE2 | 0.9922 | 0.9845 | 0.1785 | 0.2535 | 519 |
| names/HE3 | 0.9877 | 0.9756 | 0.1542 | 0.2466 | 265 |
| names/HZ2 | 0.2921 | 0.0853 | 0.3249 | 0.4492 | 43 |
| names/HZ3 | 0.2396 | 0.0574 | 0.2974 | 0.3879 | 34 |
| names/CD2 | 0.9994 | 0.9988 | 1.8970 | 2.4050 | 559 |
| names/HG | 0.7605 | 0.5784 | 0.2571 | 0.3933 | 348 |
| names/HD21 | 0.6099 | 0.3720 | 0.1767 | 0.2317 | 387 |
| names/HD22 | 0.6594 | 0.4349 | 0.1699 | 0.2232 | 387 |
| names/HD23 | 0.6513 | 0.4242 | 0.1646 | 0.2178 | 387 |
| names/CE1 | 0.8876 | 0.7879 | 3.6570 | 4.8930 | 240 |
| names/CE2 | 0.8720 | 0.7604 | 3.8760 | 4.8920 | 161 |
| names/HD1 | 0.4631 | 0.2144 | 0.2597 | 0.3491 | 351 |
| names/HE1 | 0.9070 | 0.8227 | 0.3349 | 0.4816 | 375 |
| names/HA2 | 0.4050 | 0.1640 | 0.3598 | 0.4711 | 428 |
| names/HA3 | 0.2882 | 0.0830 | 0.3131 | 0.4471 | 417 |
| names/HZ | 0.4820 | 0.2324 | 0.2967 | 0.3927 | 123 |
| names/NE2 | 0.9710 | 0.9429 | 2.1670 | 3.9190 | 136 |
| names/ND2 | 0.3097 | 0.0959 | 2.1160 | 2.7010 | 164 |
| names/NE1 | -0.0418 | 0.0017 | 1.8260 | 2.2530 | 53 |
| names/CE3 | -0.0315 | 0.0010 | 2.8640 | 3.4630 | 29 |
| names/CZ2 | -0.0507 | 0.0026 | 2.5580 | 2.9830 | 37 |
| names/CZ3 | 0.2935 | 0.0861 | 3.3860 | 4.1680 | 29 |
| names/CH2 | 0.0438 | 0.0019 | 4.2180 | 4.7470 | 35 |
| names/HH2 | 0.0501 | 0.0025 | 0.2715 | 0.3462 | 40 |

Table S3: Complete model stats on test data across classes and atom types for Model (all) Metabolome.

| title | corr-coeff | R^2 | MAE | RMSD | N |
|---|---|---|---|---|---|
| overall | 0.8723 | 0.7609 | 0.778 | 1.12 | 699 |
| overall-H | 0.8723 | 0.7609 | 0.778 | 1.12 | 699 |
| class/MB | 0.8723 | 0.7609 | 0.778 | 1.12 | 699 |
| names/H | 0.8723 | 0.7609 | 0.778 | 1.12 | 699 |
